# Time-varying Spatial Propagation of Brain Networks in fMRI data

**DOI:** 10.1101/2024.02.12.579169

**Authors:** Biozid Bostami, Noah Lewis, Oktay Agcaoglu, Jessica A. Turner, Theo van Erp, Judith M Ford, Vince Calhoun, Armin Iraji

## Abstract

Spontaneous neural activity coherently relays information across the brain. Several efforts have been made to understand how spontaneous neural activity evolves at the macro-scale level as measured by resting-state functional magnetic resonance imaging (rsfMRI). Previous studies observe the global patterns and flow of information in rsfMRI using methods such as sliding window or temporal lags. However, to our knowledge, no studies have examined spatial propagation patterns evolving with time across multiple overlapping 4D networks. Here, we propose a novel approach to study how dynamic states of the brain networks spatially propagate and evaluate whether these propagating states contain information relevant to mental illness. We implement a lagged windowed correlation approach to capture voxel-wise network-specific spatial propagation patterns in dynamic states. Results show systematic spatial state changes over time, which we confirmed are replicable across multiple scan sessions using human connectome project data. We observe networks varying in propagation speed; for example, the default mode network (DMN) propagates slowly and remains positively correlated to blood oxygenation level-dependent (BOLD) signal for 6-8 seconds, whereas the visual network propagates much quicker. We also show that summaries of network specific propagative patterns are linked to schizophrenia. More specifically, we find significant group differences in multiple dynamic parameters between schizophrenia patients and controls within four large-scale networks: default mode, temporal lobe, subcortical, and visual network. Individuals with schizophrenia spend more time in certain propagating states. In summary, this study introduces a promising general approach to exploring the spatial propagation in dynamic states of brain networks and their associated complexity and reveals novel insights into the neurobiology of schizophrenia.

## 1. Introduction

Spontaneous neural activity in the human brain can occur in the absence of external stimuli and be observed at different spatial and temporal scales [1]. The neural activity throughout the cortex has been studied via various imaging modalities, such as calcium imaging [2, 3] and voltage-sensitive dye [4]. For noninvasive, macro-scale functional neuroimaging studies, resting-state functional magnetic resonance imaging (rsfMRI) is used to understand and study intrinsic brain activity [5]. The standard approach computes temporal coupling between the blood oxygenation level-dependent (BOLD) signals, also known as functional connectivity [6]. Such studies were then extended to estimate multiple functional networks spanning the brain, including the visual and default mode networks (DMN) [7]. Among the various methodologies, independent component analysis (ICA) is a well-known approach that extracts spatial patterns of different brain networks (i.e., functional connectivity maps) and their associated temporal activity [8, 9].

Understanding the nature of spontaneous neural activity is an active area of research. Introducing temporal lags to capture whole brain lag structures and representing patterns that shift over time has been studied in the context of region of interest analyses [10]. Other approaches have focused on capturing a set of global recurring consecutive activity patterns, such as quasiperiodic patterns, highlighting propagation across areas of the task-positive and the DMN [11, 12]. Global coactivation patterns, e.g., transient neuronal coactivation patterns, can also be related to globally propagating waves [13, 14], with most focusing only on signal peaks, while other approaches focus on continuously varying contributions [15]. Other studies show the ability to capture time-varying brain connectivity, which can be helpful in diagnosing disorders such as schizophrenia [16-18]. Studies of whole-brain dynamic connectivity also focus on the temporal coupling within and between the functional domains [19, 20]. Other work has specifically evaluated spatial brain networks using a sliding window approach [21]. This work suggests that the fMRI data captures moment-to-moment voxel-wise changes within functional brain networks, such as the default mode network. However, previous studies ignore the spatial fluidity of the brain networks which evolve with time. Spatial fluidity can be defined as the transitory spatial pattern of a given functional network over time at the voxel level measurement [22].

To our knowledge, there is no work focused on quantifying propagating spatial patterns in fMRI data within spatial dynamic states (i.e., sub-states voxel-wise dynamics within specific brain networks) [23]. Here, we focus on understanding network-specific dynamic spatial state propagation. This study presents an approach to estimating multiple brain networks using ICA and capturing time-varying propagation using lagged windowed correlations and spatial dynamic state analysis. We first evaluate the replicability of the results, followed by a study focused on changes linked to mental illness. Results show clear evidence that we can detect replicable evolution of brain networks over time, such as default mode, visual, and temporal networks. Important propagation properties, such as propagation speed and pattern show variation across dynamic states. Results also show that these spatially propagating patterns are linked to mental illness. For example, studied networks showed that subjects who are diagnosed with schizophrenia have higher dwell time compared to the control group, suggesting that subjects diagnosed with schizophrenia have less activity in their brain networks and remain dominant in a particular dynamic state. This study is a first step toward understanding the complex nature of network-specific spatial dynamic propagation.

## 2. MATERIALS AND METHODS

### 2.1 Data Information and Preprocessing

In this study, two different datasets are used. The first dataset is from the Human Connectome Project for Early Psychosis (HCP-EP) [24]. The second dataset is the Functional Imaging Biomedical Informatics Research Network (FBIRN) [25]. The HCP-EP dataset is used to validate our proposed method and evaluate replicability across two scan sessions. We select a sub-section of the HCP-EP dataset where subjects are common across the two scan sessions. To evaluate the replicability of our proposed method, we used the data from the two scan sessions and matched the results after running the pipeline independently. The FBIRN dataset will be our primary dataset for evaluating links to clinical diagnosis and result analysis.

In the HCP-EP dataset, there were 163 subjects in each session remaining after preprocessing and quality control were done. The data was collected from 4 clinical recruitment sites and the consent form of each participant was collected before scanning. Medically stable male and female subjects with a confirmed psychiatric diagnosis and healthy control (HC) subjects were enrolled in the HCP-EP study. The image data were scanned using Siemens MAGNETOM Prisma 3T scanners with a multi-band sequence and a 32/64-channel head coil. The rs-fMRI data had 2 mm isotropic resolution, multi-band acceleration factor of 8, repetition time (TR) = 720 ms, and was acquired twice with posterior-anterior (PA) and anterior-posterior (AP) phase encoding. More details about the dataset can be found on the official website of the National Institutes of Health [24].

The fMRI data were preprocessed using a combination of FSL and statistical parametric mapping (SPM12) under the MATLAB 2020 environment. Before motion correction, a distortion field was calculated from the PA and AP phase-encoded field maps by the top-up/FSL algorithm to correct for intensity and geometric distortions. Then, a rigid body motion correction was performed using SPM to fix the head motions in fMRI scans. After that, the fMRI data were normalized to the standard Montreal Neurological Institute (MNI) space using an echo-planar imaging (EPI) template and slightly resampled to 3 × 3 × 3 mm isotropic voxels. The resampled fMRI images were smoothed using a Gaussian kernel with a full width at half maximum (FWHM) = 6 mm. Since dynamic functional connectivity analysis can be sensitive to the data quality, we performed quality control (QC). We selected subjects with functional data providing near-full brain successful normalization for further analysis. This yielded a total of 170 subjects. More details can be found in the following study [26].

The second dataset for evaluating links to clinical diagnosis was fBIRN (Functional Imaging Biomedical Informatics Research Network) [25]. The groups consisted of 160 typical controls with a mean age of 36.9 and 150 individuals with schizophrenia with a mean age of 37.8. There were 115 control males and 114 males with schizophrenia. Also, 45 female controls and 36 females with schizophrenia. Seven sites across the United States collected eyes-closed rsfMRI data. Consent forms were collected following the regulation of Internal review boards of the affiliated institutions prior to scanning. Six sites used the Siemens Tim Trio System, and one used the General Electric Discovery MR750 scanner. Resting-state fMRI scans were acquired following a standard gradient-echo EPI paradigm: Field of view (FOV) of 220 × 220 mm (64 × 64 matrices), TR = 2000 ms, TE = 30 ms, FA = 770º, 162 volumes, 32 sequential ascending axial slices of 4 mm thickness and 1 mm skip. Data preprocessing used a combination of various toolboxes, such as AFNI [27], SPM [28], and GIFT [29]. We used the INRIAlign toolbox to correct head motion in SPM. 3dDespike algorithm from AFNI was applied to remove outliers. Then, fMRI data were resampled to 3 mm^3^ isotropic voxels. Then, data were smoothed to 6 mm FWHM using the BlurToFWHM algorithm of the AFNI toolbox, and each voxel time course was variance normalized.

### 2.2 Independent Component Analysis

ICA is one of the most common approaches for blind source separation. It is based on the assumption that any captured signal (***x***) can be defined as a linear combination (***A***) of its latent sources (***s***), which are mutually independent such that ***x*** = ***As*** [30]. Here, ***x*** is a vector representing the captured mixtures, represents latent sources, and ***A*** is a mixing matrix where ***A*** ∈ ℝ (*N* × *M*). ICA aims to estimate an unmixing matrix called ***W*** ∈ ℝ (*N* × *M*) such that *y* = ***W****x*, which approximates the latent sources (***s***) subject to permutation and scaling ambiguities. Implementation of ICA was via the GIFT toolbox (http://trendscenter.org/software/gift). In practice, *M* > *N*, so dimension reduction is first applied. First subject-level spatial principal component analysis (PCA) was applied, and 99% of the subject-level variance was retained. Next, group-level spatial PCA was applied on concatenated subject-level principal components (PCs) for all subjects. 20 group-level PCs were selected for future analysis, which is sufficient to capture the standard large-scale resting networks. Next, Infomax ICA was applied to estimate 20 maximal ICs. Infomax was repeated 100 times, and the ICASSO framework was used to select the best (most central) component run to ensure the stability and reliability of the ICs [30]. Subject-specific ICs and associated time courses were derived using spatially constrained ICA, group information guided-ICA using the group map as the reference [31]. Finally, different brain networks, such as the DMN, visual, temporal lobe, and subcortical network were identified based on their spatial maps and power spectra [32].

### 2.3 Calculating Spatial Maps with Lagged Window Correlation

The dynamic spatial propagation of various brain networks can be evaluated at a voxel-wise level. For this purpose, the temporal coupling between the selected brain network and every brain voxel using the lagged sliding window approach was calculated. Same cleaning procedures were followed, effectively capturing dynamic patterns on time courses and every brain voxel to reduce noise [33]. The cleaning procedure included orthogonalizing to estimated subject motion parameters, linear detrending, despiking, and bandpass filtering using a fifth-order Butterworth filter (0.01–0.15 Hz). The tapered window was obtained by convolving a rectangle (width = 30 TRs: with a Gaussian (σ = 3 TRs), and the sliding step size of 2 TR was used. In this study, a parameter, lag (τ) was introduced, to indicate the temporal correlation between a given brain network and every voxel of the brain for every (*t* ± τ).

As the sliding window shifts along the time course of the target network, temporal correlation between the windowed time points located at time *t* of the selected ICA time course and the windowed time points located at (*t* ± τ) for every value of τ is calculated. In more general terms, multiple voxel-level temporal correlations by shifting the window over bold time signals for different τ values are computed while keeping the ICA network’s window location fixed.

For example, if τ = 2 and TR = 2 sec is selected, then a set of 5 spatial maps located at (-4 sec, -2 sec, 0 sec, 2 sec, 4 sec) for every sliding step *t* will be captured. These are called ‘lag points’. More generally, if τ = *n*, the 2*n* + 1 spatial maps are computed for each window step. Each window’s correlation variation at different times was captured by introducing the lag parameter.

### 2.4 Calculation of Dynamic Spatial States

After calculating the lagged spatial maps for every subject, k-means clustering was used to identify the spatial dynamic states and associated patterns. For this study, the cluster number of 4 was selected, estimated by the elbow method. This was replicated 50 times with different initializations using the k-means++ method to increase the probability of avoiding local minima [34]. The correlation distance was used as the metric to calculate the similarity between the data points. The analysis of dynamic states was done in two steps. First, ran k-means clustering for each subject and captured exemplar (or reference) states. Next, run a second level of k-mean, merging all the exemplar states to obtain exemplar centroids. These centroids are used as a reference for calculating the final centroids, computed by concatenating all subject’s spatial maps and running k means clustering. The complete pipeline is shown in Figure 1

**Figure 1:**
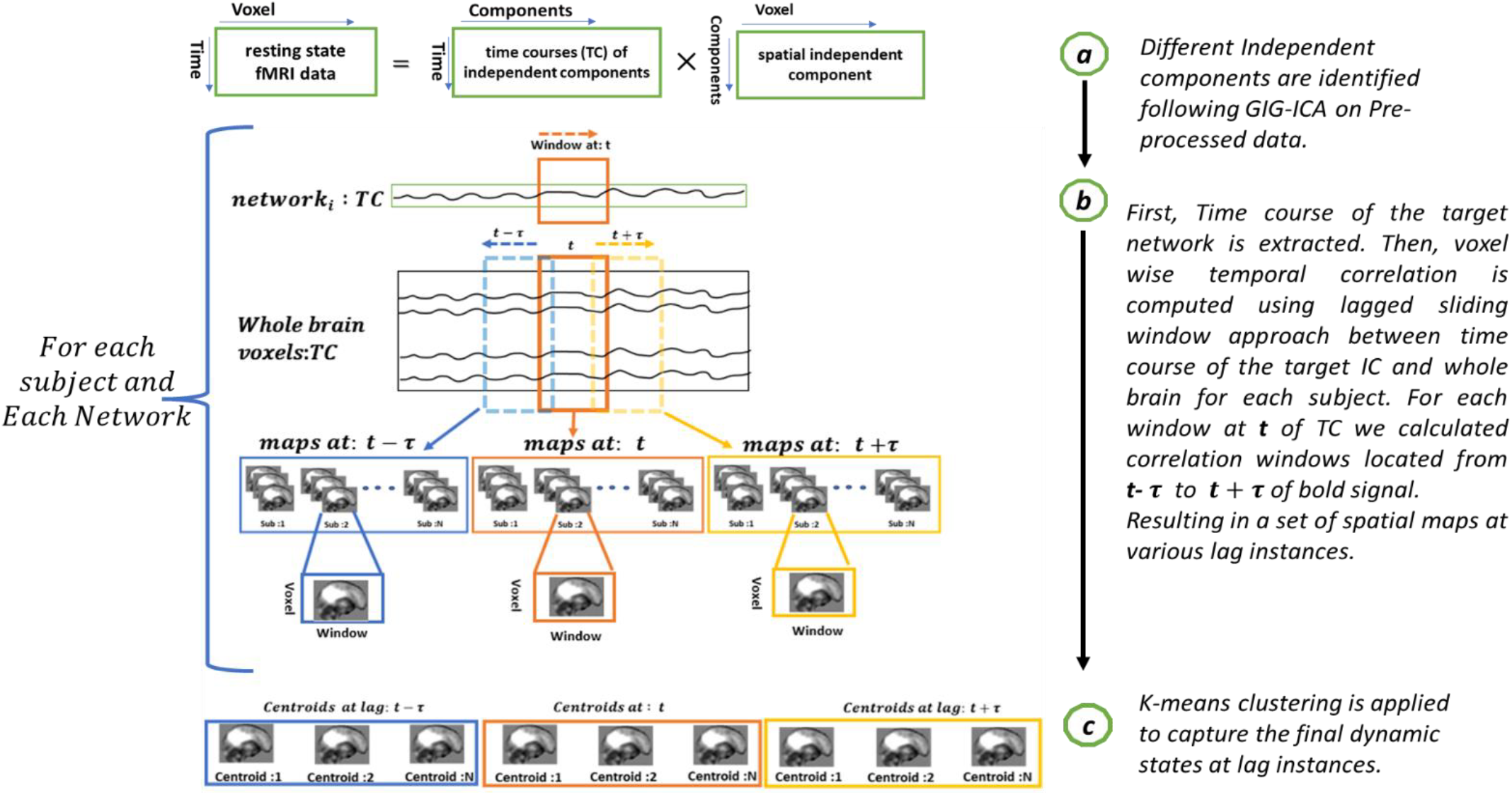
Analysis Pipeline of the proposed lagged windowed correlation approach. The proposed pipeline uses a lagged sliding window approach to capture voxel level network propagation that evolves with time.

### 2.5 Statistical Experiments

A number of summary measures were calculated on the output, including mean dwell time (MDT), fraction time (FT), and disperse rate (DR). Mean dwell time calculates the average time spent at each state before transitioning to other states. Fraction time gives the percentage of total time in each state. Disperse rate represents the similarity between spatial maps at different lag points, which tells us how fast the network propagates. We observed group differences based on the calculated features.

## 3. EXPERIMENTS AND RESULTS

In a data-driven approach, replicability is an important factors both for validity and interpretability [35]. Replicability ensures that given same subjects with different data produce similar results. The proposed method was tested over two sessions of the HCP-EP dataset to evaluate this. Analysis was run over the same subjects over two different sessions, and the final aggregated centroids, called dynamic states, in two independent analyses. After obtaining two sets of dynamic states, correlation was calculated between the centroids in each session and between sessions. The results show that the dynamic state captured by the two sessions was highly correlated. Figure 2 shows highly replicable results across multiple sessions. Figure 2(a) represents the spatial similarity between spatial maps in the dynamic states calculated from data using session 1. Cell values represent how similar the spatial maps across the states are calculated from session 1 data only.

**Figure 2:**
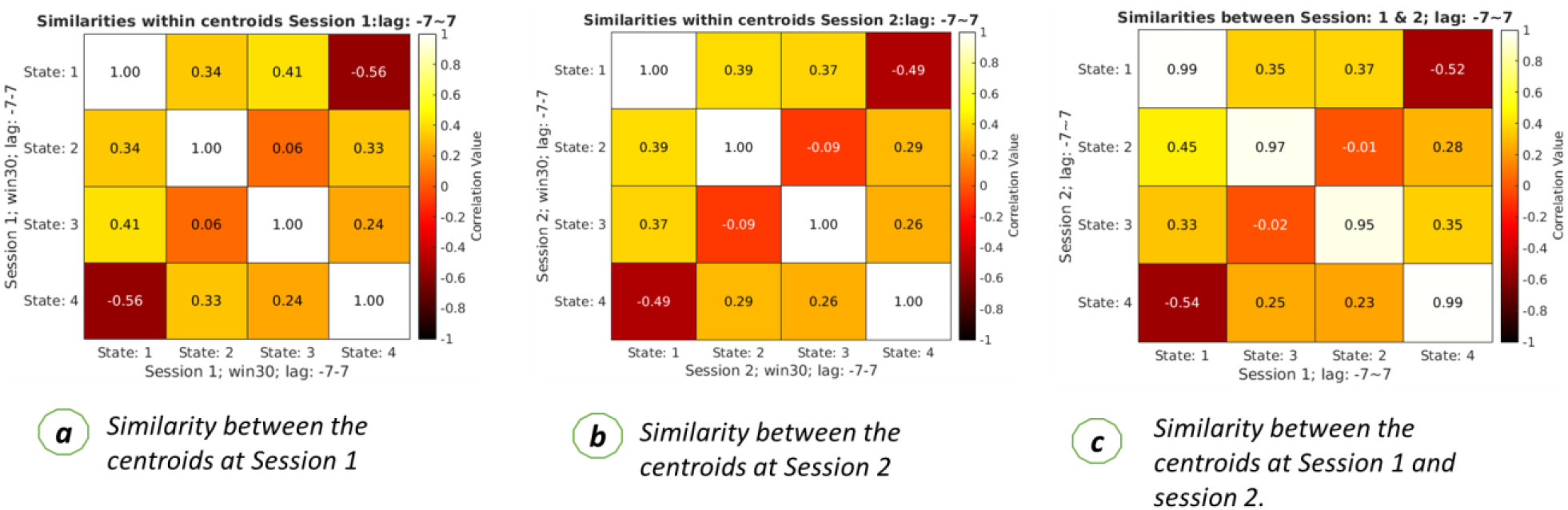
Replicability of results across multiple sessions. The left figure represents the spatial similarity between the centroids or dynamic states calculated from session 1 data. Each cell represents the correlation coefficient between two centroids calculated from clustering. The middle figure represents the spatial similarity between the dynamic states or clustering centroids calculated from session 2 data. The correct figure represents the spatial similarity between the centroids of session 1 and session 2 data. This shows a high similarity between the states between the two sessions, which can be seen from the main diagonal cells. The high correlation value proves that the proposed pipeline produces reproducible centroids over multiple sessions.

Similarly, Figure 2(b) represents the spatial similarity between the spatial maps at different states computed from session 2 data. Figure 2(c) represents spatial similarity between the spatial maps located at session 1 against the spatial maps of different states of session 2. After reordering the states based on the maximum similarity, it was observed that spatial maps at state 1 of session 1 and session 2 were highly similar as they had a high correlation value of 0.99. A similar observation was observed in the case of state 4 of both session 1 and session 2. Also, spatial maps at state 2 of session 1 were highly similar to spatial maps calculated at state 3 of session 2 with correlation values of 0.95-0.99. The primary diagonal value of Figure 2 (c) shows how the spatial maps at different states are similar. These results also provide evidence that the proposed pipeline provides replicable results.

Next, to observe the spatial propagation of brain networks, we plotted the dynamic spatial maps of the networks. In Figure 3, we plotted the default mode network. With our proposed approach, we were able to capture the spatially propagating patterns over time. Figure 3 shows how the dynamic states of the DMN propagate in different states. We also can observe the DMN network remains positively correlated for a certain period before it slowly becomes anti-correlated.

**Figure 3:**
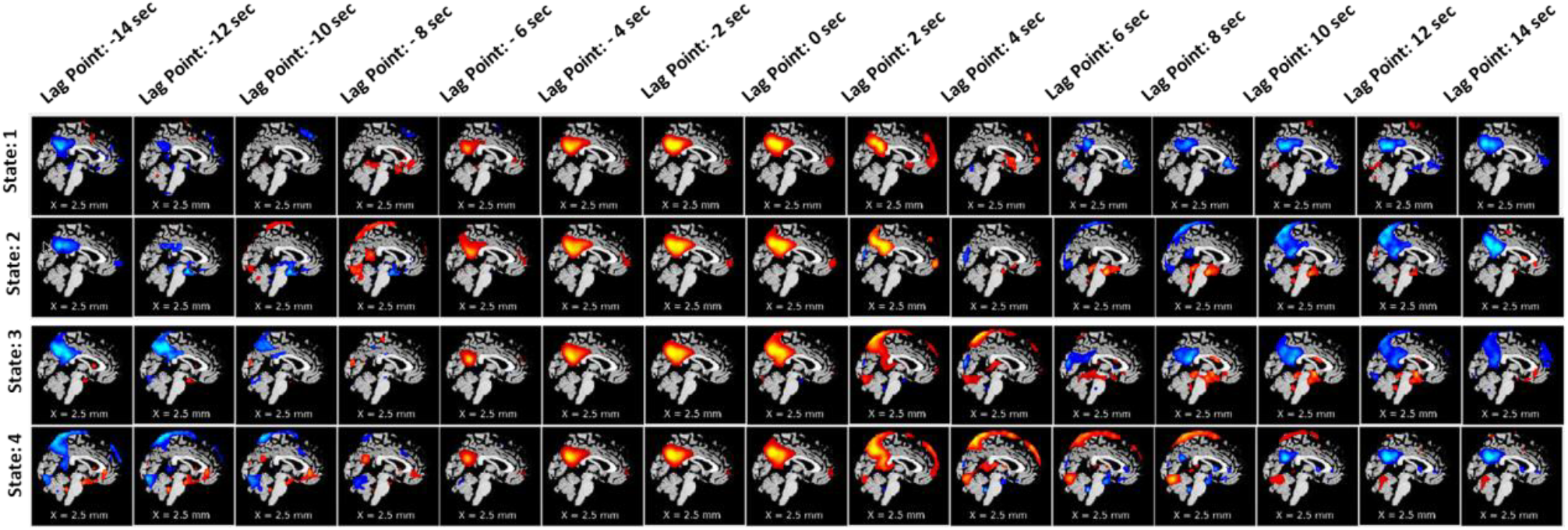
spatial propagation of the DMN over time. The figure shows how different dynamic states of DMN are fully formed early at a time (lag point: -4 sec) and slowly dissipate at lag point 0 seconds (state 3). Also, the speed of disperses varies from state to state. It also shows that DMN stays positively correlated with the BOLD signal for around 6-8 second (starting from the lag point: -6 second to lag point: 2 seconds) before it slowly dissolves and become anti-correlated.

To observe the spatial propagation of other networks, visual, subcortical, and temporal were selected. The spatial propagation patterns were observed after running the pipeline independently for each network. Figure 4 shows the visual network, Figure 5 shows the sub-cortical network, and Figure 6 shows the temporal network. From the figures, it is observed that different networks propagate similarly. However, the propagation rate is different for each network individually.

**Figure 4:**
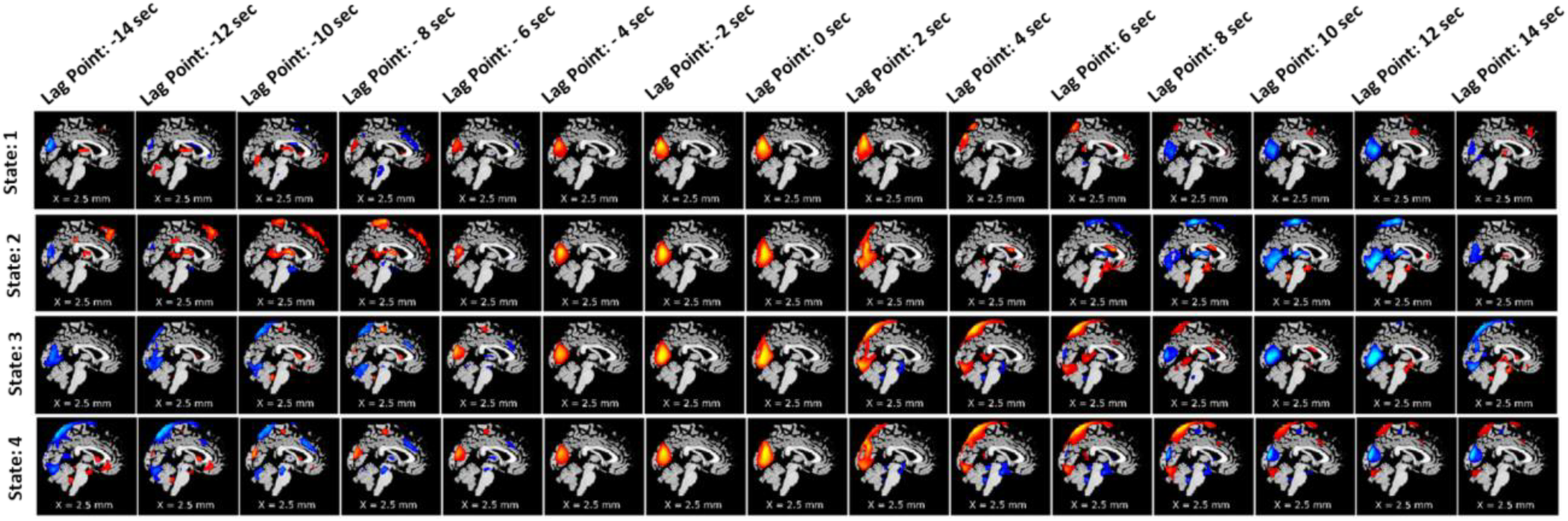
Spatial propagation of the visual network over time. The figure shows how different dynamic states of the visual network are fully formed early (lag point: -4 sec) and start to dissipate at lag point 0 seconds. Also, the speed of disperses varies from state to state. It also shows that the visual network stays positively correlated with the BOLD signal for around 4 seconds (starting from the lag point: -4 second to lag point: 0 seconds) before it slowly dissolves and become anti-correlated.

**Figure 5:**
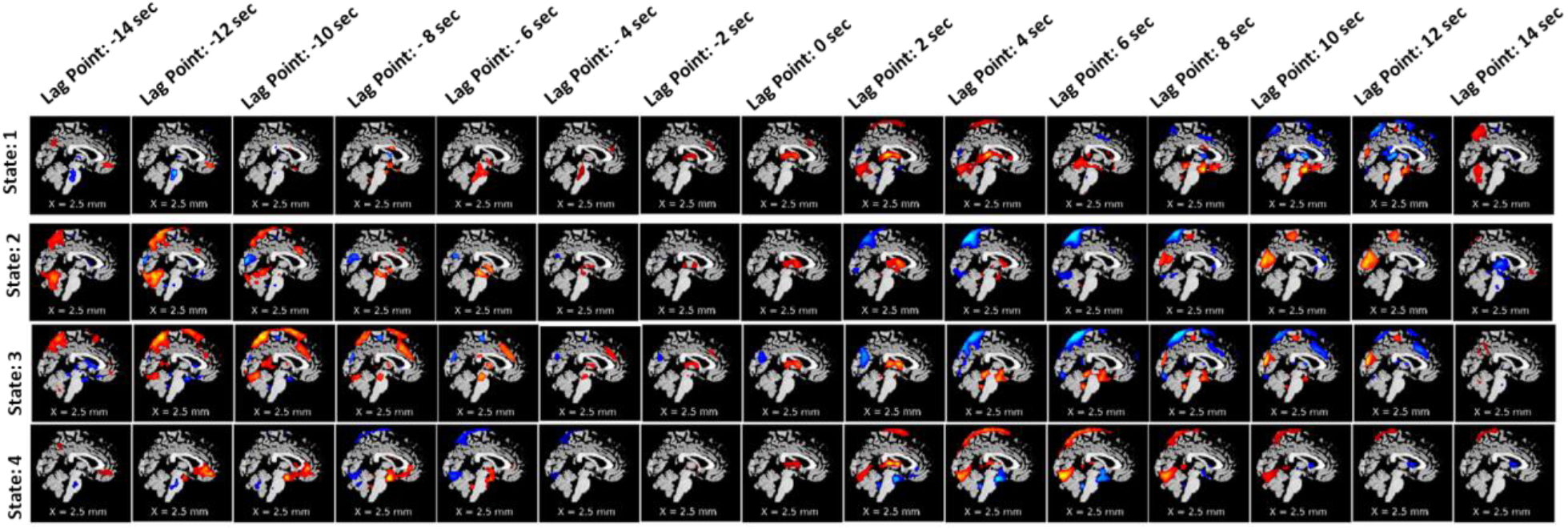
Spatial propagation of the sub-cortical network over time. The figure shows how different dynamic states of the sub-cortical network are fully formed at the time (lag point: 0 sec) and start to dissipate at lag point 2 seconds. Also, the speed of disperses varies from state to state. It also shows that the sub-cortical network stays positively correlated with the BOLD signal for around 2 seconds (starting from the lag point: 0 seconds to lag point: 2 seconds) before it slowly dissolves and becomes anti-correlated.

**Figure 6:**
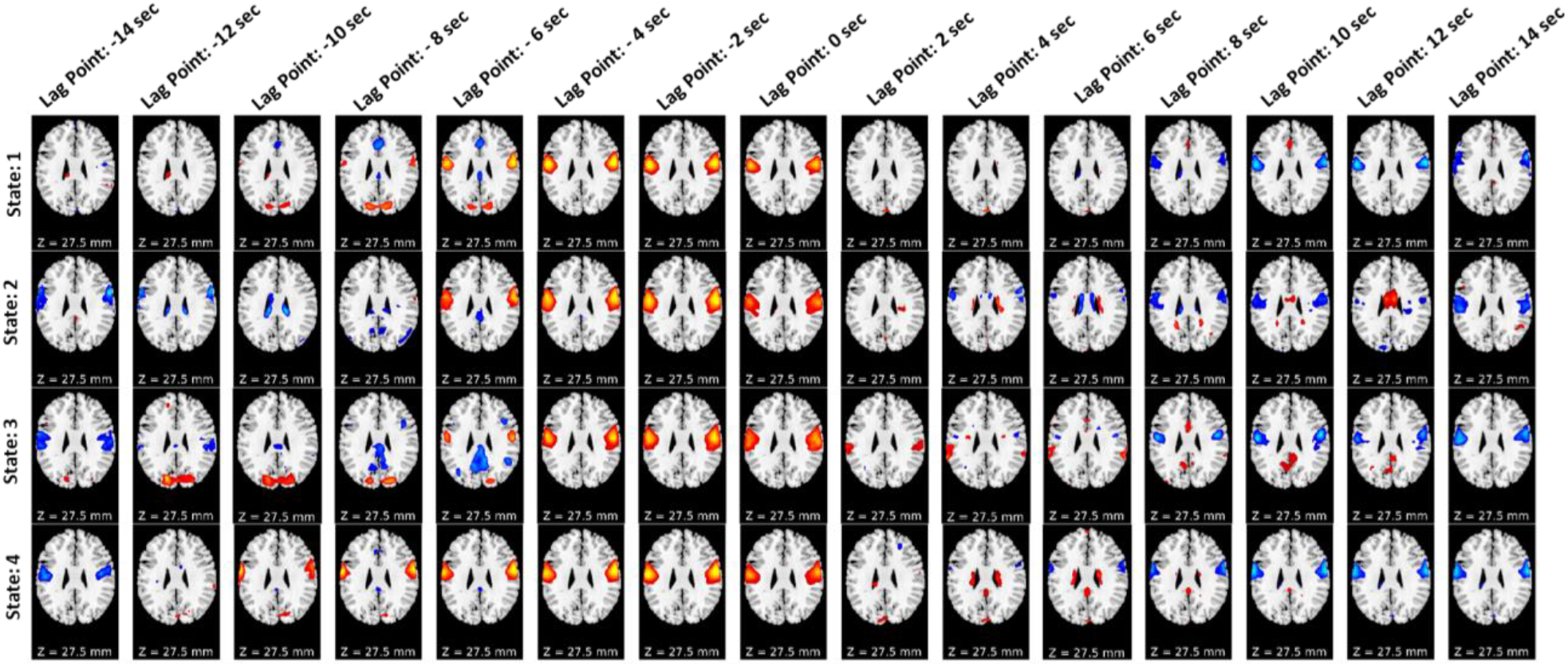
Spatial propagation of the temporal network over time. The figure shows how different dynamic states of temporal network are fully formed early at the time (lag point: -4 sec) and start to dissipate quickly at lag point 2 seconds. Also, the speed of disperses varies from state to state. It also shows that the temporal network stays positively correlated with the BOLD signal for around 4 seconds (starting from the lag point: -4 second to lag point: 0 seconds) before it quickly dissolves and become anti-correlated.

To analyze the importance of various spatial patterns that were captured, several measurements, such as mean dwell time (MDT), were calculated for each spatial map. First, the group differences based on the MDT at different lags were computed and plotted. It captured group differences at different lag points for different states. In a lag-less analysis, this information is never captured. Figure 7-10 plots the average difference of MDT of two groups, and significant states are marked. For example, we observed group differences at State 2, State 3, and State 4 when the lag was at 8 seconds in DMN. However, only state 2 and state 4 shows significant group difference at lag 0 seconds.

**Figure 7:**
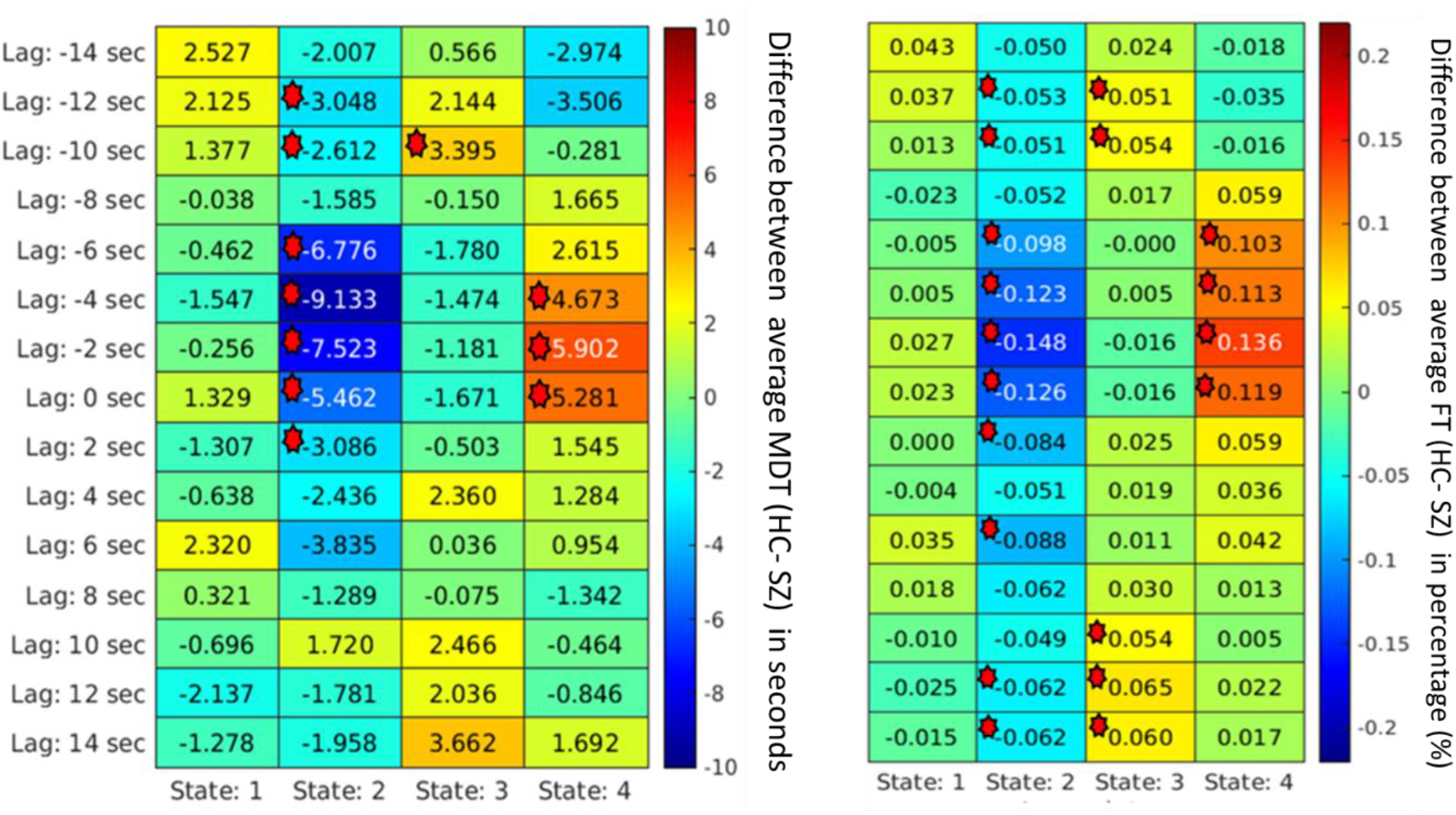
*For the visual network*, the mean dwell time between two groups captures significant differences at different states at different lags. Each cell value represents the difference between the two groups’ average MDT (HC-SZ). Significant differences (p < 0.05 (FDR corrected) are marked. State 2 and state 4 captured significant group differences where in state 2, most schizophrenia subjects dwelled longer; in state 4, healthy controls dwelled longer on average in visual network states where schizophrenia patients dwelled more captured significant group differences.

**Figure 8:**
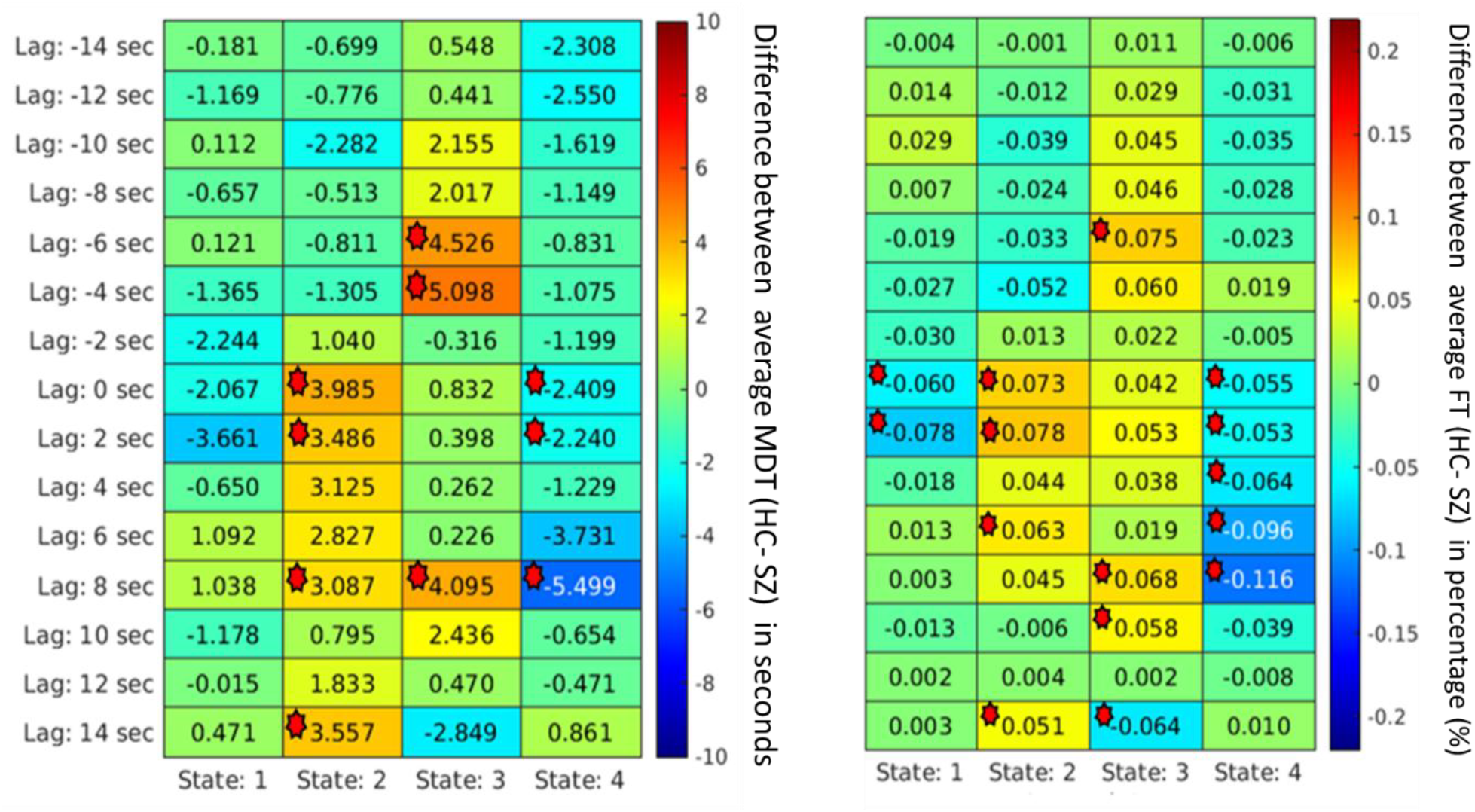
*For the default mode network*, Mean Dwell Time between two groups captures significant group differences at different states at different lags. Each cell value represents the difference between the two groups’ average MDT (HC-SZ). Significant differences (p < 0.05) are marked. The Schizophrenia group dwells more in state 1 and state 4 than the healthy controls, unlike visual network states where healthy control resides more capture significant group difference.

Similarly, we can capture group differences for all the states if we observe lag at -8 sec and -2 sec for the subcortical network shown in Figure 9. Similarly, the FT was computed to capture group differences between the two groups. Figure 7 – 10 plotted the FT calculated at different lag points for all dynamic states. It was observed that FT at other lag points changes with time as the network propagates with time, and FT changes, too. However, different states located at different lag points capture significant group differences.

**Figure 9:**
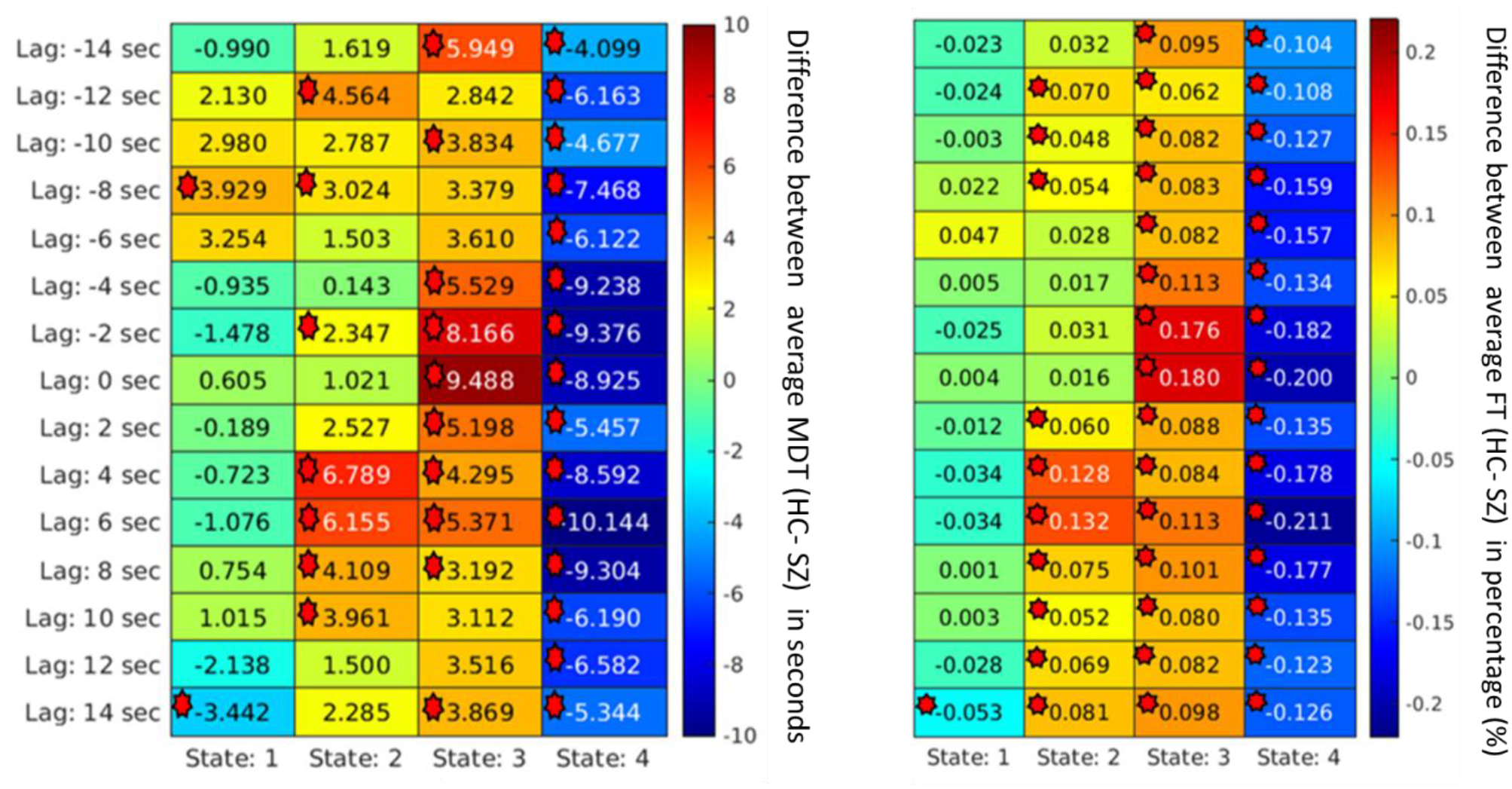
*For the subcortical network*, the Mean Dwell Time between two groups captures significant group differences at different states at different lags. Each cell value represents the difference between the two groups’ average MDT (HC-SZ). Significant differences (p < 0.05) are marked. Most of the schizophrenia group dwelled in most of the time in state 4 over the evolving time. On average, most healthy controls also resided in state 3, suggesting groups do not change conditions between the dynamic and subcortical states.

**Figure 10:**
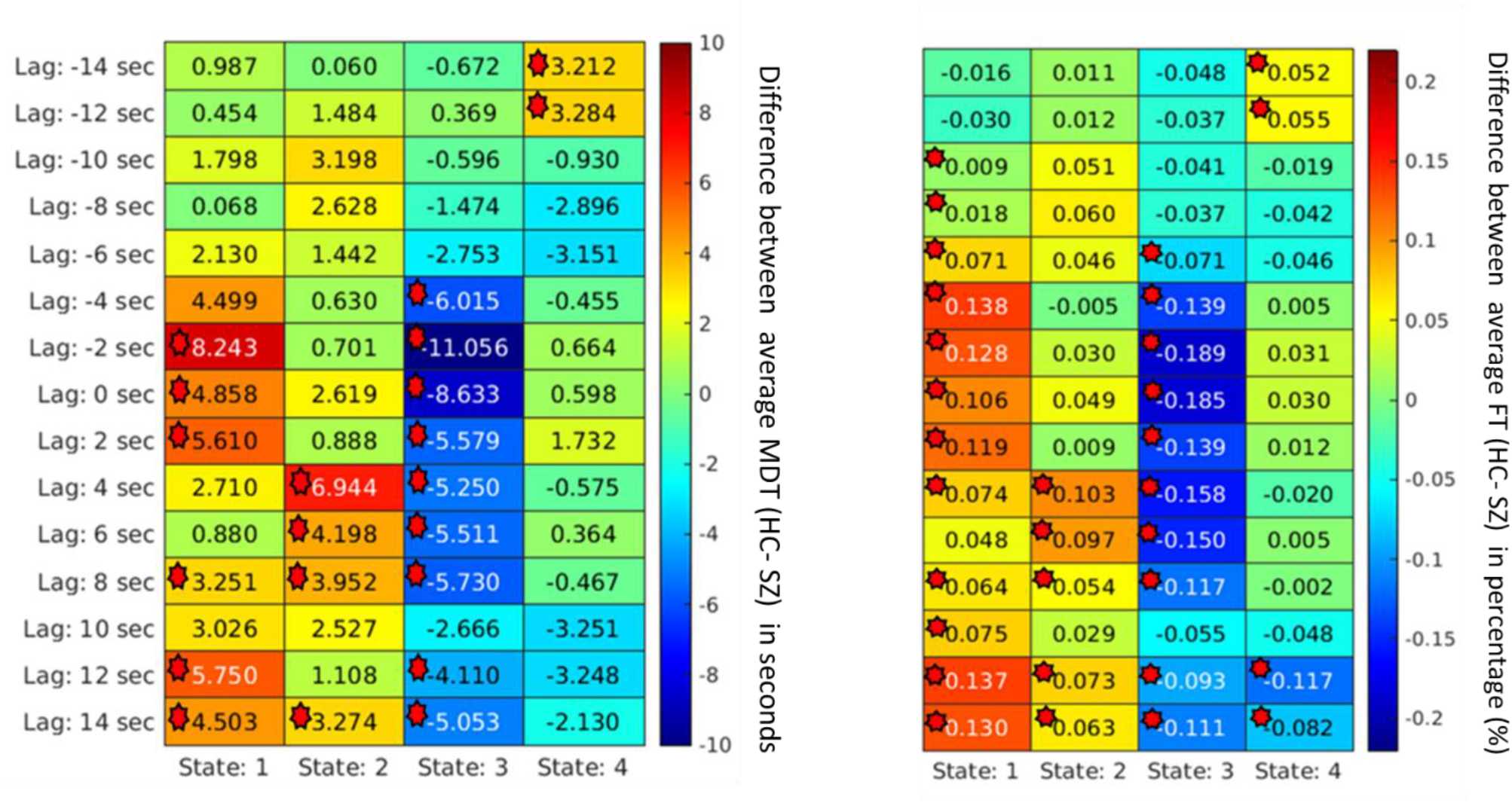
*For the temporal network*, the Mean Dwell Time between two groups captures significant group differences at different states at different lags. Each cell value represents the difference between the two groups’ average MDT (HC-SZ). Significant differences (p < 0.05) are marked. After the network is formed, the schizophrenia group dwells more in a particular state, such as state 3, and healthy control resides more in state 1 most of the time.

Although some significant states are not shown in the figure due to p-value correction, figure 7-10 captured an overview of different dynamic states, capturing information regarding the group differences related to neurological illness. Results reveal some interesting patterns among the spatial states, such that most group effects are located around the center when the network is more visible, and lag is minimal. Also, it was observed that in some dynamic states, subjects with schizophrenia dwell significantly longer than the healthy control group on average.

## 4. Discussion

The brain is a complex dynamic system. Static analysis of fMRI data may provide important information; however, such research cannot capture the time-varying behavior of the brain. Recent studies have proposed various ways to capture important aspects of brain dynamism by brain connectivity. Still, these studies ignore the brain’s spatial aspects, which evolve with time. The proposed study focused on how each brain network propagates with time over space. The proposed approach showed that spatial information of brain networks changes with time. This study examined four brain networks individually and found that spatial pattern changes as the BOLD signal propagates across the brain. The spatial patterns may differ based on the selected network. However, the observed brain network showed similar propagation behavior. Observing these propagating patterns was the primary goal of this study. There has been little work in this area focused on understanding the dynamics of the propagation of time-varying states of the brain.

We observed that the rate of the propagation of a network also differs based on the starting state of that network. The correlation between the spatial maps at a particular state can give the propagation rate. This supports our primary hypothesis of the presence of network-level propagation.

In neuroimaging dynamic analysis, replicability is a considerable concern. In our proposed pipeline, strict correspondence cannot be enforced between different runs due to the data-driven nature of the approach. For example, ICA may not always produce the same network in the same order. Calculated dynamic states occur in different orders. Despite these challenges, we show that the proposed analysis produces similar states across multiple runs, irrespective of their ordering, in data collected from the same subjects at different times. That supports the hypothesis that the proposed analysis has a high level of replicability.

This study captures how the brain network patterns propagate over time. As the signal travels through the brain, the brain networks interact. From our analysis, we captured dynamic spatial maps and their propagation patterns with respect to time. Each map captures the spatiotemporal correlation values between the ICA time course and the BOLD signals. The maps also represent which brain parts are related to the target network that becomes active with time. For example, from the spatial map propagation maps, we can observe that at different lag points, different brain areas show activation other than the target network, which suggests that brain networks are related and interact with each other as signals travel through time.

We also observed that the ICA network becomes positively correlated and more visible with time. Most of the significant group differences were captured within such periods. However, in some cases, we also observed that while the network can capture group differences in different dynamic states, it remains visible, although anti-correlated, with respect to the BOLD signal. This suggests that anti-correlated spatial patterns can also capture important neurological information related to illness. Such anti-correlated brain network patterns were captured before in traditional dynamic studies, suggesting that different networks are anti-correlated with respect to temporal correlations [36, 37]. Our study shows that these anti-correlated patterns are time dependent. Also, these anti-correlated patterns suggest that the networks do not work independently; instead, different networks share a time-dependent correlation between them. Previous static analysis did not capture such information.

We also observed that subjects with schizophrenia dwell in a particular state longer before they traverse to the next state. For example, state 3 of temporal, state 3 of visual, state 4 of DMN, and subcortical shows the dwell time of subjects with schizophrenia is larger than that of healthy control subjects. From the state maps, we also observed how different sub-networks become active with time. For example, different sub-networks of the ICA subcortical network were active at different lag points, and each of them was able to capture significant group differences based on calculated MDT. Because we had a model order of 20, which is considered a lower model order, some of the subnetwork overlapped, and these networks do capture group differences when they become active with time. This explains why we were able to capture group differences when the target network was not very visible.

We also observed that each network has a particular state where schizophrenia group members dwell longer than other states. These states also were identified as showing a significant group difference. Networks such as the temporal lobe, visual, and subcortical showed a similar pattern. Previous studies also observed that dwell time in one of the dynamic states was longer than other dynamic states [36, 37], which is aligned with our observations.

Furthermore, our results show that lag-based analysis can capture more states than lag-less analysis results where no lag is used. These additional lagged states can also capture substantial group differences and convey potentially important and distinct information. Also, it was observed that different states contain different amounts of information. For example, MDT and FT between groups change as the spatial pattern varies with time. This also strengthens our second hypothesis that spatial patterns at different times contain significant information that can be useful for identifying groups and clinical purposes. Such states were ignored in previous studies.

## 5. Limitations and Future directions

The results presented in this study have several primary methodological and experimental limitations. We only presented four brain networks in the current study. Also, our analysis ignores the presence of inter-network interaction. In future studies, we will explore these inter-network relationships and how networks relay information. We will also extend the work to extract more features that would help us understand the complexity of spatial propagation patterns.

## 6. Conclusion

This study proposed a novel approach to observe the propagation patterns in the dynamic spatial states of brain networks. The proposed method was tested against data collected from two sessions of the HCP dataset, which showed high replicability. The study focuses on the evolution of spatial patterns of various brain networks that vary over time, which have not been studied previously. These time-varying spatial networks are statistically significant for group analysis between patients with schizophrenia and controls. In summary, this study can be considered a building block toward thoroughly understanding the complexity of global propagation.

